# Intra-population variability in genome-wide repressive histone marks underpins differential gene expression in a fungal wheat pathogen

**DOI:** 10.1101/2025.09.19.677456

**Authors:** Leen Nanchira Abraham, Ana Margarida Sampaio, Suhani Bhattacharyya, Sabina Moser Tralamazza, Daniel Croll

**Author notes:** Co-first authors (Leen Nanchira Abraham, Ana Margarida Sampaio).

## Abstract

Epigenetic modifications influence the expression of phenotypic traits by modulating gene expression and responses to environmental cues. In plant pathogens, the expression of virulence-associated genes such as effectors and gene clusters encoding the production of secondary metabolites are known to be regulated by epigenetic modifications. Modulating epigenetic patterns of such genes is considered a key adaptation for pathogens to successfully attack hosts. Gene expression variation within pathogen species are regulated by extensive *cis-* regulatory polymorphism and insertion activities of transposable elements. However, whether pathogens vary in epigenetic profiles among members of the same species remains largely unexplored. Here, we focus on the major fungal wheat pathogen *Zymoseptoria tritici* and establish histone methylation profiles for 45 isolates of an extensively characterized wheat field population. We analyzed the facultative heterochromatin mark H3K27me3, a histone methylation that is thought to regulate effector and gene cluster loci in the genome. H3K27m3 coverage was increased in transposable element rich regions, with newly inserted long-terminal repeat retrotransposons contributing to epigenetic variation among pathogen genotypes. Overall, nearly 20% of all genes showed within-population variation in H3K27me3 marks, which likely contributes to the substantial within-population variation in gene expression. Effector candidate genes and members of gene clusters showed higher than average variation in repressive histone marks among isolates. Taken together, our study provides among the first insights into intra-species epigenetic variation of a fungal pathogen. Such population-level variation in histone methylation patterns opens avenues to recapitulate epigenetic mechanisms of pathogen adaptation.

## Introduction

Phenotypic variation among individuals may arise from mutations or non-genetic changes (1). Major epigenetic processes, including DNA methylation, histone modification, and various RNA-mediated processes regulate spatial and temporal gene expression patterns (2). Among these, covalent modifications of histones modify the local chromatin structure and affect DNA accessibility to transcription factors regulating gene expression (2). Chromatin is organized by the nucleosome, an octamer with 147 base pairs of DNA wrapped around histone proteins. Histone proteins (H2A, H2B, H3, and H4) are made up of a globular domain and an unstructured tail domain that can be modified by acetylation, methylation, phosphorylation, and ubiquitylation (3). Among these, methylation of histone tails is well studied and has diverse important biological functions (4). Methylation of histone H3 at lysine 4 residue (H3K4me) is a broadly found mark for transcriptionally active and gene-dense regions in eukaryotes. Although many aspects of H3K4me mechanisms and functions appear to be shared among kingdoms, there are significant differences between H3K4me and transcription in plants and mammals. For instance, plants appear to lack preferential co-localization of H3K4me3 and H3K27 as has been shown in mammals. Similarly, H3K4me2/3 and DNA methylation seem to be mutually exclusive in *Arabidopsis thaliana* (5). Methylation of histone H3 at lysine 9 residue (H3K9me) is typically found in repeat-rich regions near transposable elements and satellite repeats causing transcriptional silencing (6). In addition to heterochromatin formation, H3K9 methylation is a prerequisite for DNA methylation, another type of epigenetic modification that is involved in gene silencing (7,8). Comparative analyses of DNA methylation in eukaryotes showed that budding and fission yeasts are devoid of DNA methylation (9). *Saccharomyces cerevisiae* also lacks repressive histone H3K9 methylation suggesting yeast lineages lost this epigenetic pathway (10). The trimethylation of histone H3 at lysine 27 (H3K27) modification underpins facultative heterochromatin and is associated with regions of responsive gene expression regulation (11). Unlike H3K9 methylation which prevents the binding of transcription factors and causes a persistent state of silencing, H3K27 allows genes to be activated through transcription factor binding in response to environmental cues and stress (3,12,13). Despite extensive knowledge of gene functions and domains preferentially associated with different chromatin states, how chromosomal regions gain specific marks remains poorly understood.

Histone methylation marks are not conserved within species and epigenetic variation can underpin gene expression variation (14). In *A. thaliana*, H3K27 occupancy varies little, and the flanking transposable elements appear to account for most variation (15). In humans, histone tail modifications are highly variable but stably inherited across generations (16). Hundreds of sequence variants in the human genome are identified to be associated with both histone modification and gene expression variation consistent with widespread epigenetic effects on gene expression (17). DNA binding molecules that bind to specific DNA motifs can recruit or stabilize histone modifications (18). In humans and rats, histone-associated DNA motifs have shown significant overlap with the expression of quantitative trait loci SNPs, suggesting an important role in gene regulation (19). Histone methylation levels vary in rat genomes in response to *cis*- and *trans-*acting regulatory factors, indicating that histone trimethylation marks are impacted by genetic variation (20). Insights from these studies strongly suggest that epigenetic variation in histone methylation marks are widespread within species and likely affect adaptive phenotypic trait variation. However, studies on the extent and consequences of histone methylation polymorphism in natural populations are lacking with few exceptions (21).

In fungi, epigenetic modifications can facilitate the invasion of their hosts (22). Plant pathogens need to rapidly respond to environmental cues when in contact with a plant host (23). Plants produce a variety of molecules to inhibit fungal development (24). In turn, fungal pathogens secrete secondary metabolites and small proteins (*i.e.* effectors) to manipulate the host (25). Up-regulation of pathogen genes upon infection is highly concerted and characterized by an initial wave of effector genes and specialized metabolite gene cluster expression (26–32). Regulatory control of some effector genes is governed by epigenetic changes related to H3K9me3 marks such as in the rapeseed pathogen *Leptosphaeria maculans* (33). Effector genes residing in similarly repeat-rich and repressive regions in the wheat pathogen *Zymoseptoria tritici* are upregulated by a reduction in H3K9me3 and H3K27 (34). Infection-related metabolite gene clusters are largely regulated epigenetically with frequent associations with H3K27 in the fungal pathogens *Epichloë festucae, Fusarium graminearum, F. fujikuroi and Colletotrichum higginsianum* (35–38). Histone methylation plays also broader roles in the evolution of fungi with H3K27 marks underpinning reduced transcriptional robustness (39) and the loss of H3K27 contributing to elevated mutation rates (40).

The wheat pathogen *Z. tritici* causes severe yield losses under conducive conditions and has spread globally with the introduction of the host (41). Gene regulation is governed by numerous expression quantitative trait loci (eQTLs) located close to transcription start sites (42). The genome is organized into distinct compartments of gene-dense regions of open chromatin and repeat-rich regions with repressive H3K9me3 and H3K27 marks (43). TEs actively create new copies reshaping the genomic landscape, impacting gene expression, phenotypic trait variation, and genome size (44–48). The species has lost a functional DNA methylation machinery in its recent evolutionary history though (49). H3K9me3 and H3K27 histone methylation marks affect negatively and positively the base mutation rate, respectively (40). Strikingly, H3K9me3 supports and H3K27 reduces the stability of degenerated accessory chromosomes (50). The species carries vast polymorphism at the genetic and transcriptional levels (29,41,42,47,51–53). Gene regulation is governed by *cis* and *trans-*acting loci with most genes showing evidence for at least one regulatory region (42). However, structural variation and movements of TEs likely impact also the epigenetic landscape (44–47) with consequences for the expression of individual genes.

Here, we generated genome-wide H3K27me3 profiles using chromatin immunoprecipitation sequencing (ChIP-seq) for a highly polymorphic population of 45 *Z. tritici* isolates. The experiments were conducted in a minimum culture medium simulating early phase of infection. We first searched for genomic factors underpinning H3K27me3 variation among genotypes such as gene density and recent TE activity. Then, we assessed the impact of H3K27me3 variation on gene expression and analyzed variation in histone methylation marks across gene functions encoded by the pathogen genome. We focused in particular on epigenetic variation near effector genes and secondary metabolite gene clusters.

## Results

### Population chromatin immunoprecipitation sequencing (ChIP-seq)

We performed genome-wide ChIP-seq analyses of H3K27m3 marks in a highly diverse population of *Z. tritici* (*n* = 45) isolated from a naturally infected Swiss wheat field (53) (Fig 1A). Isolates were propagated individually under standardized, nutrient-deprived culture conditions mimicking early plant infection stages (44). We obtained between 339-985 Mb of sequencing data per isolate. The mapping rate of reads against the reference genome IPO323 range from 76.3% to 92.1% comparable to mapping rates for whole genome sequencing data of the same population (47) (Supplementary Table S1, Supplementary Figure S1A). Between 54.2% to 76.7% of mapped reads were uniquely aligned to the reference genome (Supplementary Figure S1A). After duplicate filtering, we retained between 7 - 20 million mapped reads per isolate. We calculated the normalized strand cross-correlation coefficient (NSC) as an indicator of the H3K27 signal-to-background noise ratio. The NSC was consistently above 1.05 indicating adequate enrichment in H3K27m3 signals (Supplementary Figure S1B). We assessed read distribution biases across the genome and found background uniformity (Bu) values to be mainly above 0.8 indicating low false positive peak calling risks (Supplementary Figure S1C). The peak calling on the aligned H3K27m3 ChIP-seq reads produced between 1571-9428 peaks (average 2953) among isolates (Supplementary Figure S1D) stemming from an average of 56% of all aligned reads (FRiP score; Supplementary Figure S1E).

**Figure 1:**
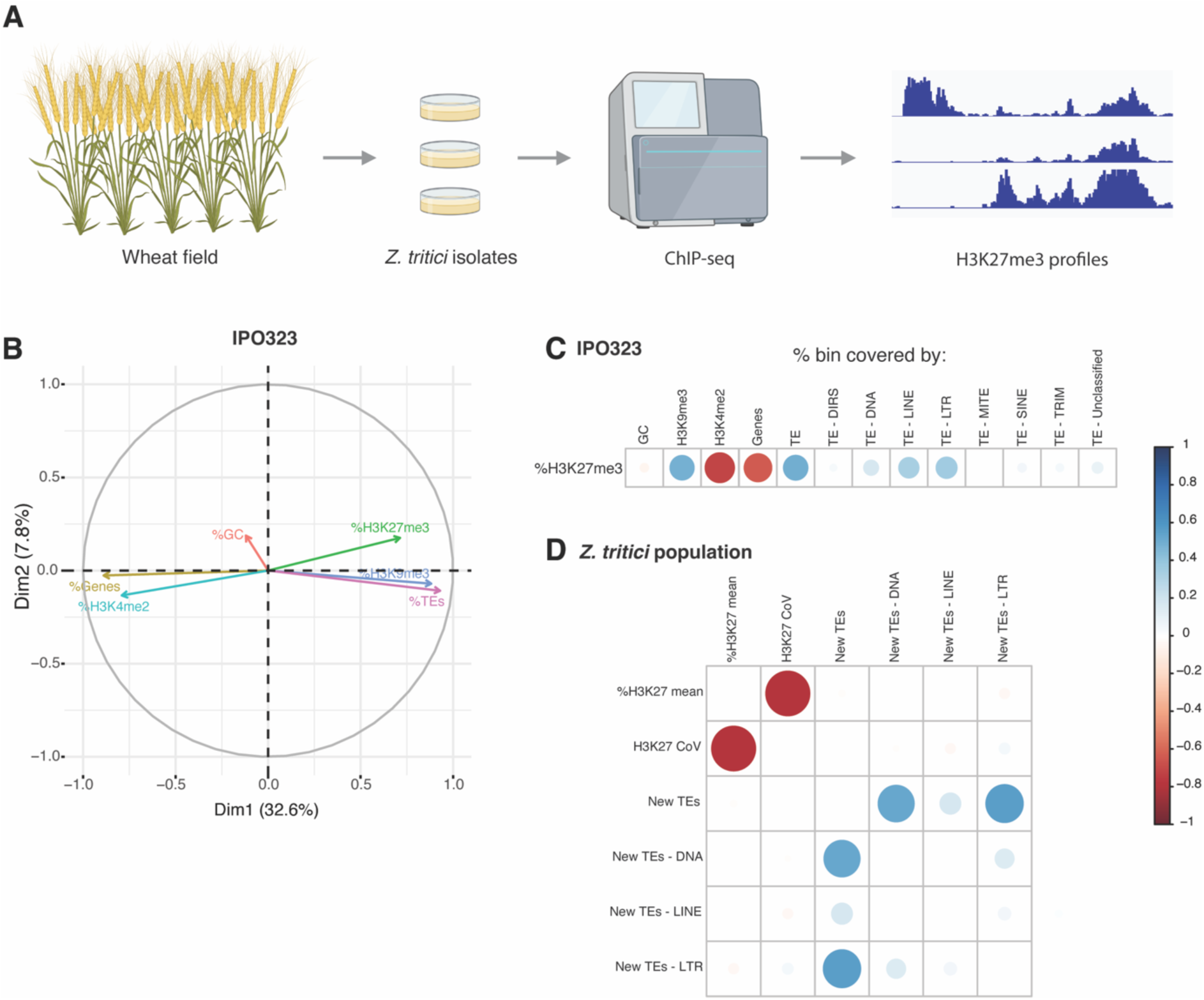
Experimental design and genomic features correlated with histone methylation mark variation. A) Schematic overview of the study design. B) Ordination plot of GC content, H3K27m3, H3K9m3, H3K4m3 coverage, coding sequence density, and transposable element (TE) density in the reference genome. The metrics were assessed in genomic windows of 10 kb. C) Correlation plot between H3K27m3 coverage and GC content, H3K9m3, H3K4m2, coding sequences and TEs (incl. superfamily categories) in the reference genome. D) Correlation plot between mean H3K27m3 coverage and the coefficient of variation among bins, as well as counts of recently inserted TEs (incl. superfamily categories) among isolates of the analyzed population.

### Genomic factors associated with histone methylation variation

To assess genomic factors associated with histone methylation variation among isolates, we first explored ChIP-seq peak distribution in the reference genome using 10 kb windows. For the first analyses, we focused on previously generated ChIP-seq data for the reference genome isolate IPO323 and the marks H3K27m3, H3K9me3 and H3K4me2 (54). GC content ranged from 31-58% with an average of 52% among windows (Supplementary Table S2). TEs varied widely (0-100%) among windows with an average occupancy of 19%, while coding sequences occupied on average 43% of the windows (Supplementary Table S2). The active histone mark H3K4me2, typically associated with promoters and enhancers of active genes, was the most frequently found mark across the genome consistent with the high gene density. H3K4me2 was positively correlated with coding sequences, while repressive histone marks (H3K9me3) were positively correlated with regions rich in TEs (Fig 1B), especially DNA transposons, LINEs and LTR retrotransposons (Fig 1C, Supplementary Figure 2A). Since the presence of TEs was positively correlated with H3K27m3, we assessed whether the most recent TE activity (*i.e.* new insertions) was associated with variation in H3K27m3 variation among isolates. Using the newly generated ChIP-seq population survey for H3K27m3, we defined the H3K27m3 mean coverage and coefficient of variance (CoV) for each genomic window using data for all 45 field isolates. Regions of high H3K27m3 among isolates were positively correlated with newly inserted LTR retrotransposons (Fig 1D). Topologically associated domains (TADs) described for the reference genome IPO323 were shown to be enriched in H3K27m3, in particular on accessory chromosomes (55). Here, we observed that individual TADs located in both core and accessory chromosomes exhibit also variation in H3K27me3 coverage among isolates (Supplementary Figure 2B).

### Within-population variation in H3K27m3 gene body coverage

We found substantial variation among isolates in the proportion of H3K27m3 marked genes (Fig 2A). Marks were consistently higher for genes on accessory chromosomes compared to core chromosomes (Fig 2B and C). Gene body coverage showed a strongly bimodal distribution of either no or complete H3K27m3 coverage (Fig 2D; Supplementary Figure S3A). We found that 69.2% and 11.4% of the genes showed either consistent H3K27m3 coverage or no coverage among isolates, respectively. The remaining 19.4% of the genes showed variable gene body coverage by H3K27m3 marks within the population (Fig 2E, Supplementary Table S3).

**Figure 2:**
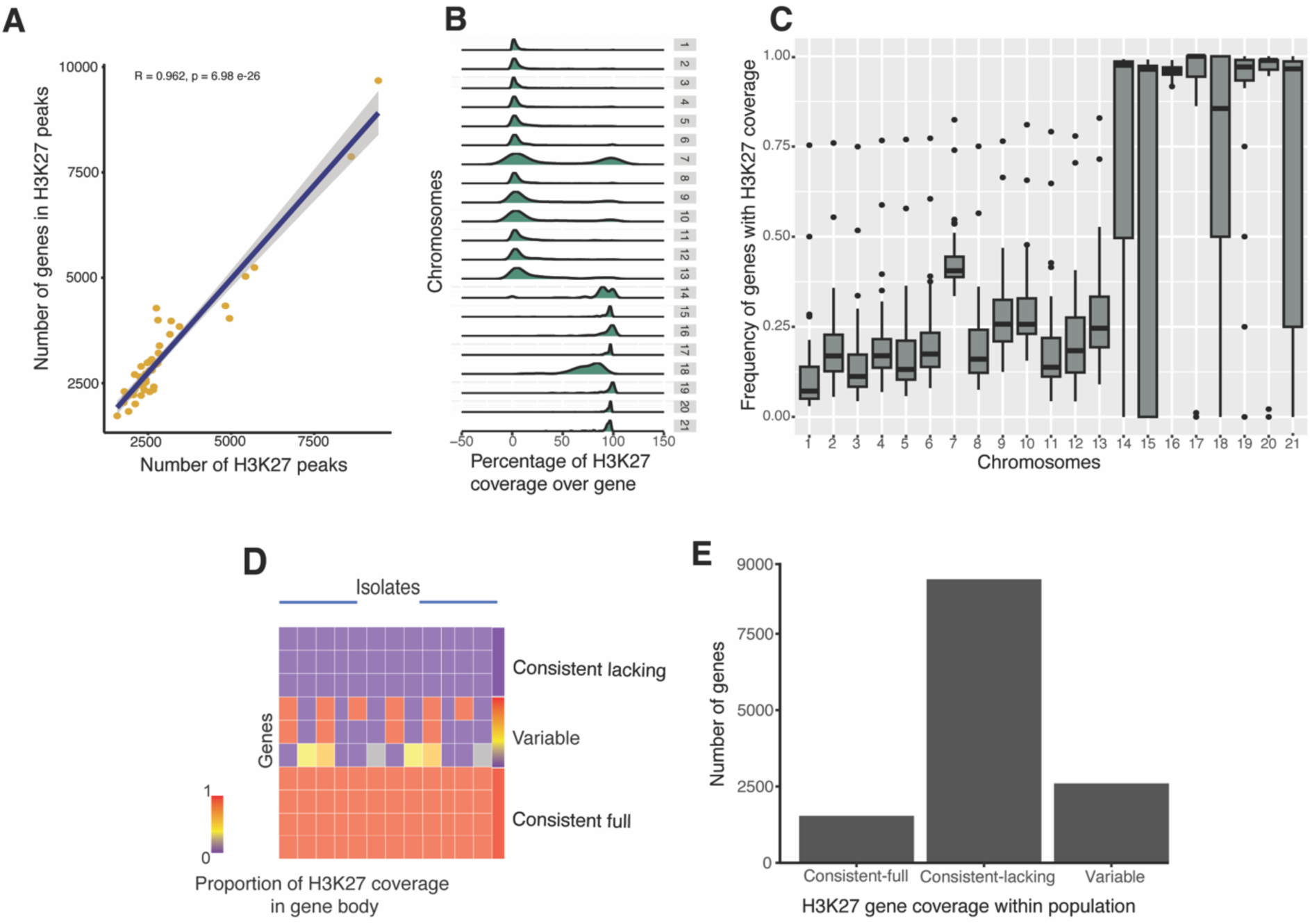
Genome wide variation in H3K27m3 methylation marks covering gene bodies. (A) Number of genome-wide genes overlapping with H3K27m3 peaks. (B) Average profile of H3K27m3 coverage density across gene bodies throughout chromosomes. (C) Density of H3K27m3 coverage across gene bodies on chromosomes within the population. (D) Illustration of the different gene categories based on H3K27m3 coverage in the population. (E) Number of genes in each gene category defined by H3K27m3 coverage.

### Epigenomic variation of pathogenesis-related genes and gene clusters in the population

We analyzed H3K27m3 mark variation in gene bodies for two important categories of virulence-associated functions: candidate effector genes and gene clusters encoding secondary metabolite biosynthetic pathways. We found that most of the candidate effector genes (56.2%) showed a consistent lack in H3K27m3 coverage across the gene body, while 13.4% exhibited consistent complete coverage, and 30.4% showed variation in H3K27m3 gene body coverage among isolates (Fig 3A). By analyzing the expression variation of candidate effector genes under the same nutrient-poor conditions used to obtain H3K27m3 profiles, we observed that expression variation was higher for effector genes consistently covered by H3K27 marks (Fig 3B). However, no significant differences in expression variation have been found comparing effector genes either consistently covered, consistently lacking, or with variable H3K27m3 coverage (Fig 3B), suggesting that this facultative repressive marker might not act as a decisive factor for expression under the tested conditions. We investigated the epigenetic profile of the major effector gene encoding *AvrStb6*, a small, secreted effector triggering resistance responses in wheat cultivars carrying the *Stb6* gene in a gene-for-gene (GFG) interaction (56). The large majority (82%) of the isolates showed complete coverage by H3K27m3 marks (Supplementary Table S4). The absence of H3K27m3 marks is more consistently explained by the absence of the *AvrStb6* itself, rather than the existence of unmarked variants of the effector gene (Supplementary Figure S4A). Loss of *AvrStb6* was shown to occur at a low frequency in various populations (57). Consistent coverage by H3K27m3 is tied to silencing of *AvrStb6*, but lack of H3K27m3 coverage was associated with both high and low *AvrStb6* expression (Fig 3C).

**Figure 3:**
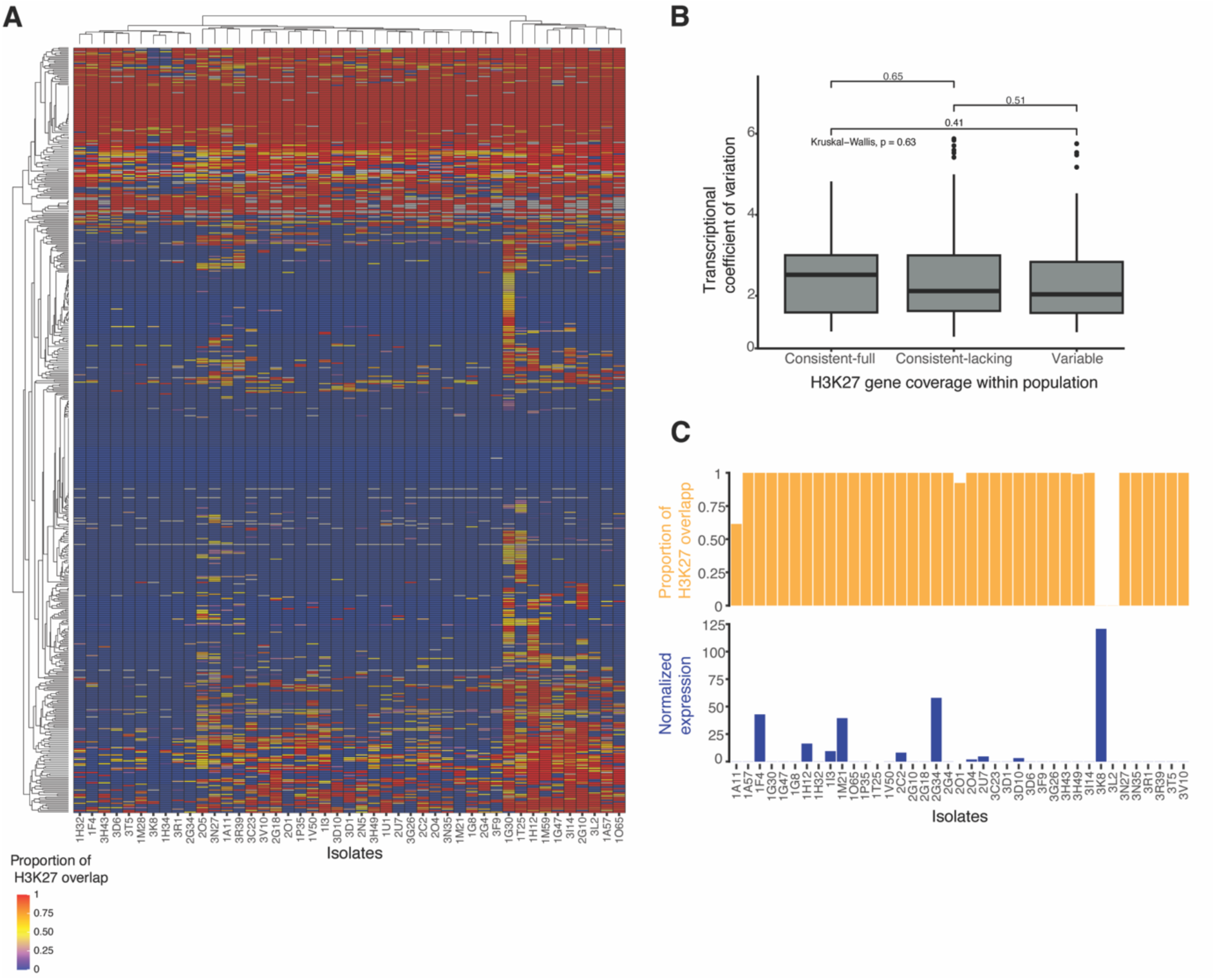
H3K27m3 gene body coverage variation among candidate effector genes. A) H3K27m3 profile in *Z. tritici* effector candidate genes. B) Transcriptional variation of effector genes across gene categories defined by H3K27m3 coverage within the population. C) *AvrStb6* H3K27m3 coverage and expression per isolate.

We also investigated five candidate effector genes (58), which were previously shown to vary in their expression among isolates during infection (29). These include genes encoding a hydrophobin (ZtIPO323_12724), a DNase (_6487), a ribonuclease (_7456) cytotoxic against plants and microbes, a putative phytotoxin inducing necrosis and defense responses in plane trees (_8099) and gene of unknown function (_4110). Most genes showed variation in H3K27m3 coverage among isolates (Supplementary Figure S4B), consistent with the observed expression variation. The only exception was ZtIPO323_6487 with consistent gene body coverage by H3K27m3 among isolates (Supplementary Figure S4B).

To investigate variation in H3K27m3 gene body coverage of secondary metabolite gene clusters, we analyzed 788 genes forming 39 predicted gene clusters (Supplementary Table S5). We found that 59.5% of all gene clusters consistently lack H3K27m3 marks among isolates, 24.9% of genes showed variable H3K27m3 coverage, and 15.5% were consistently covered by H3K27m3. Genes encoding (core and additional) biosynthesis, regulatory, or transport functions exhibited similar levels of variation in H3K27m3 mark coverage, ranging between 20.7 and 30% among isolates (Fig 4A). Most of the secondary biosynthetic gene clusters exhibited different H3K27m3 coverage patterns. In contrast, genes encoding terpene synthesis clusters (26 genes in four clusters) were consistently lacking H3K27m3 marks (Fig 4B). Biosynthetic core genes showed also high variation in H3K27m3 coverage among isolates (Fig 4C). For instance, a polyketide synthase (PKS) gene cluster shows variation H3K27m3 mark variation in both the biosynthetic and additional biosynthetic genes of the cluster (Fig 4D). A PKS core gene (ZtIPO323_4401), which has been associated with antimicrobial activity against other fungi was also showing a highly variable H3K27m3 profile (59,60) (Fig 4D). We observed no variation in H3K27m3 marks for a biosynthetic core gene link to abscisic acid production (ZtIPO323_2921) experiencing strong upregulation during the transition from the biotrophic to the necrotrophic phase (Fig. 4E) (52).

**Figure 4:**
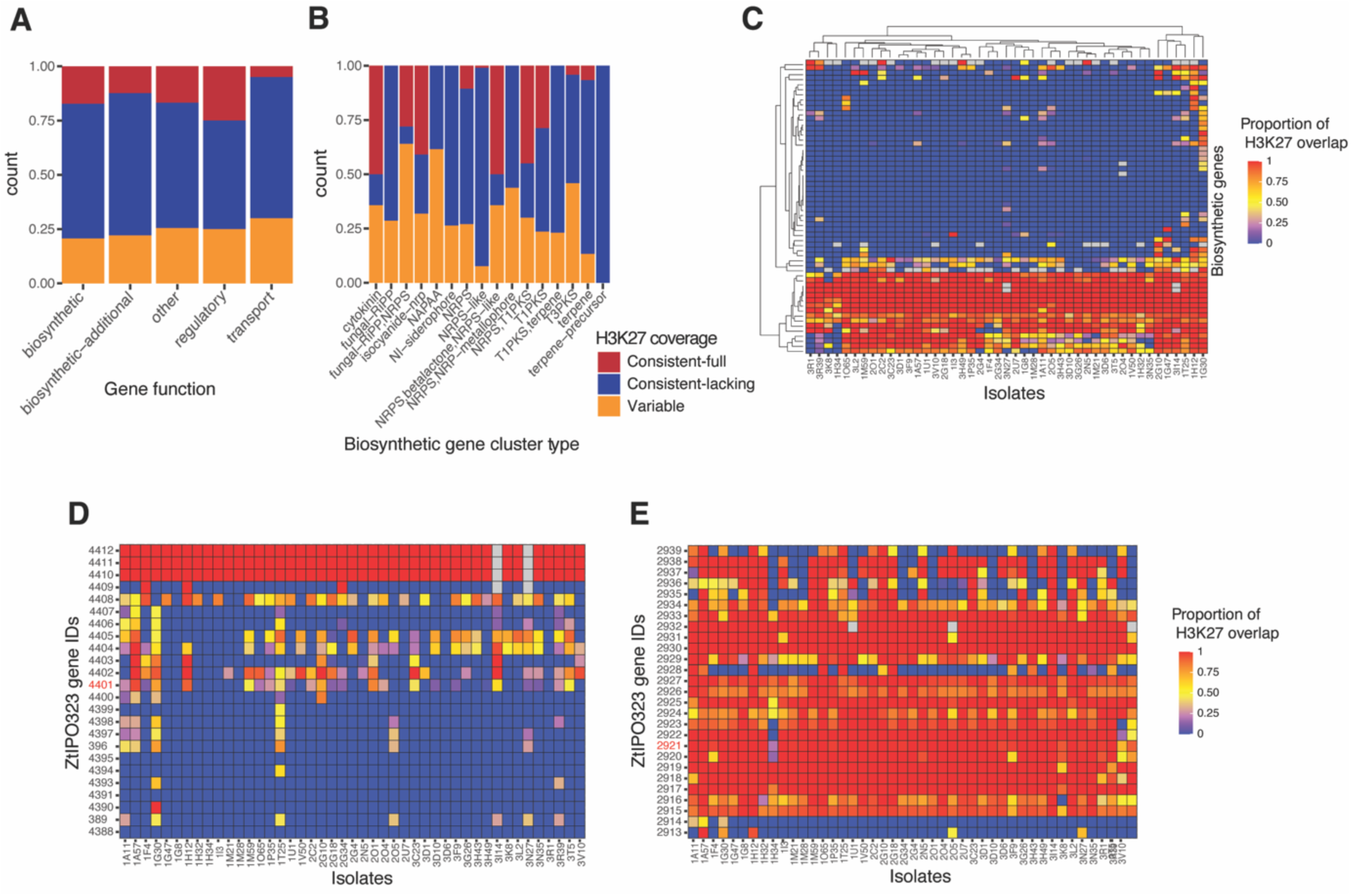
H3K27m3 gene body coverage of secondary metabolite gene clusters. (A) H3K27m3 coverage profile by secondary metabolite gene functions. (B) H3K27m3 coverage profile by biosynthetic gene cluster type. (C) H3K27m3 profile of secondary metabolite core biosynthetic genes. (D) H3K27m3 profile of abscisic acid–encoding biosynthetic gene cluster. Core biosynthetic gene is marked in red. (E) H3K27m3 profile of polyketide synthase–encoding biosynthetic gene cluster. Core biosynthetic gene is marked in red.

## Discussion

We produced the first genome-wide assessment of population-scale variability in repressive histone modifications in a fungal plant pathogen. Our analyses show that gene presence and the activity of TEs are correlated to H3K27m3 marks across the genome. We also observed that the proportion of H3K27m3-marked genes vary among members of the same species and that genes on accessory chromosomes typically show higher proportions of marked genes. Approximately one fifth of all genes show heritable variation in H3K27m3 marks even in a single population of the pathogen. Interestingly, we also found a large-scale repressive histone mark variation for genes important for pathogenicity (*i.e.* effector candidates and gene clusters involved in the biosynthesis of secondary metabolite).

Perturbations in the chromatin state of a chromosomal region can mediate transcriptional variation among individuals (16). TEs are often marked by repressive marks such as H3K9me3, acting as a powerful repressor of their transcription and transposition activity (61,62). However, H3K27m3 is also positively correlated with increased TE transcription both in *Z. tritici* and the rice blast fungus *Magnaporthe oryzae* (32,63). In *M. oryzae*, H3K27m3 was found broadly enriched across TE-rich regions, especially near effector genes (32). In *Z. tritici*, H3K27m3 marks are more abundant particularly near DNA transposons, LINEs and LTR retrotransposons. In *M. oryzae,* LINEs and LTR retrotransposons showed also H3K27me3 marks but now for DNA elements (32). LTRs have been broadly found to be targeted by PRC2, responsible for catalyzing H3K27me3 (64). Here, we show that regions with LTRs are indeed associated with H3K27m3 variation within the species suggesting that the LTR targeting may be heterogeneous and possibly dynamic given the ongoing activity by LTRs within the species (48).

Genes can also be marked by facultative repressive histone markers. Our within-species analyses showed that a substantial minority of all genes exhibited variation in H3K27me3 mark coverage among isolates, representing a substantial epigenomic variability. Considering the transcriptional silencing associated with H3K27m3 marks over the gene body (65), the observed H3K27m3 variability likely plays a role in gene expression variation observed within the species (42). Consistent with (43),we also observed a striking difference in H3K27m3 marks between core and accessory chromosomes. Genes located on accessory chromosomes were found to have a high proportion of H3K27m3 marks consistent with their likely dispensable roles. Furthermore, in *Z. tritici*, the loss of H3K27m3 increases the stability of some accessory chromosomes suggesting an important role of H3K27m3 in genome integrity (50). Among core chromosomes, genes located on chromosome 7 showed comparatively higher H3K27m3 overlap with the gene body. This observation may be linked to the proposed origin of the right arm of chr 7 from an accessory chromosome (43). Moreover, chromosome 7 showed an enrichment of TEs compared to other core chromosomes. Flanking TEs may spread the repressive chromatin marks to the neighboring gene and downregulate gene expression (33,44). The low proportion of regulatory variants mapped for genes on chromosome 7 is consistent with the proposed role H3K27m3 plays on accessory chromosomes (42). Our study revealed intra-species variation in H3K27m3, suggesting that such epigenetic differences may contribute to the observed variation in gene expression across the population.

Histone modifications were widely studied in the context of pathogenicity-associated genes such as effector-encoding and secondary metabolite genes (66,67). The epigenetic variation associated with pathogenicity-related genes within species could constitute adaptive genetic variation selectable for more optimal responses to environmental cues. We observed a high variability of H3K27m3 marks for candidate effector genes (30.4%) and secondary metabolite encoding genes (24.9%) in the population compared to background genes (19.4%). H3K27m3 also governs gene regulation as a response to various environmental stress factors. For example, in the rice blast fungus *M. oryzae,* effector genes marked by H3K27m3 during axenic growth were affected by chromatin dynamics and transcriptional variation during host infection (32). The associations of H3K27m3 with genes encoding proteinaceous and metabolic effectors or proteins involved in stress responses are known also from the fungal plant pathogen *Leptosphaeria maculans* (33). The role of H3K27m3 in the regulation of secondary metabolite gene clusters in fungi has been also extensively studied (68,69). Here we observed that genes encoding biosynthesis (core and additional), regulatory, transport functions and other exhibited similar levels of variation in H3K27 marks, ranging between 20.7 and 30%. This suggests that a local and histone-mediated regulation of such genes, apart from the transcription factors mediating global regulation of metabolite clusters (68). We observed variable H3K27m3 profiles in the biosynthetic genes of the PKS secondary metabolite gene cluster (ZtIPO323_4401). The presence of a SNP located in the 3’UTR of this same gene, has been previously associated with an antimicrobial active at least against the basidiomycete *Albatrellus confluence* (59,60). Hence, histone methylation profiles and analyses of polymorphism associated with trait expression can be combined.

Further studies to understand the genetic basis of the H3K27m3 variation within species will provide a picture of the complex interplay of genetic and epigenetic variation in the PKS encoding secondary metabolite gene cluster. Our study established the first population-level histone methylation profile for a fungal species. We found extensive variation in H3K27m3 marks spanning a large part of the gene body. The epigenetic variation present within a single field population highlights the challenge to contain rapidly evolving pathogens.

## Materials and methods

### Chromatin immune-precipitation and sequencing

Isolates of *Z. tritici* (*n* = 45) were collected from an experimental wheat field planted with different cultivars in 2016 (53,70). Isolates were grown for 10 days in a modified Vogel’s (Minimal) Medium N with ammonium nitrate replaced by potassium nitrate and ammonium phosphate, without sucrose and agarose to induce hyphal growth (71). We performed chromatin immune precipitation on the collected mycelia following (54): 2.5 mL of 20% formaldehyde (final concentration ∼0.5%) added directly to the flask and incubated for 15 min at room temperature while shaking (100 rpm). Formaldehyde was quenched by adding 2 mL of 2.5 M glycine followed by centrifugation at 2000 rpm for 5 min and the pellet was washed with 1 X PBS. The resulting pellet (150mg) was frozen in liquid nitrogen and homogenized using mortar and pestle. Ice-cold lysis or chromatin buffer was added in a ratio of ∼5 µL chromatin buffer to 1 mg of pellet. Micrococcal nuclease (MNase, #M02479; NEB) was added to the reaction and incubated for 10-20 min in a 37°C water bath and mixed every 2 min by inversion. To stop the reaction, 4 µL of 0.5 M Na-EGTA (pH 8.0) was added and the samples were placed on ice. Samples were mixed and centrifuged at 4000 rpm for 5 min at 4°C. Then 800 µL supernatant was transferred to a fresh tube and non-specific proteins were pre-cleared with magnetic Dynabeads (Invitrogen) by incubation at 4°C on a rotator for 1h. Then, samples were centrifuged at 5000 rpm for 1 min. The histone H3K27me3 antibody (pAb) from Active Motif (Cat. No. 39055) was added (5 µL) to each sample tube and incubated overnight at 4°C on a rotator. After the overnight incubation, 20 µl of magnetic Dynabeads were added and incubated for 2 hours at 4°C on a rotator to allow for antibody binding. Samples were placed on a magnetic rack and the supernatant was discarded. The pellet was washed with 1 mL of cold ChIP lysis buffer, 1 mL of ice-cold LiCl, and 1 mL of ice-cold TE buffer. Next, 63 µL of 65°C TE buffer with Sodium do-decyl sulfate was added and incubated for 10 min at 65°C, resulting in the elution of DNA from the beads. The elution was repeated with another 63 µL of warm TES and supernatants were pooled. Chromatin crosslinks were reversed by incubating samples for 6-16 h in a 65°C incubator. To the de-crosslinked samples, 125 µL of water and 1.9 µL of 20 mg/mL RNase A were added, and tubes were incubated at 50°C for 2 h. Subsequently, 9.5 µL of 20 mg/ml proteinase K was added and tubes were incubated for another 2 h at 50°C. The resulting supernatant was cleaned and eluted in 30 µL of nuclease-free water using Wizard® SV Gel and PCR Clean-Up System. The ChiP-sequencing library was prepared for sequencing and analyzed using a NovaSeq 6000 in paired-end mode with a read length of 150 bp.

### Quality control and peak calling

Raw ChIP-seq sequencing data was checked for quality using FastQC v0.12.1 (72) and trimmed with Trimmomatic v0.39 (73) to remove adapter sequences and low-quality reads based on the following parameters: ILLUMINACLIP: TruSeq3-PE.fa:2:30:10 LEADING:3 TRAILING:3 SLIDING WINDOW:4:15 MINLEN:36. Trimmed sequences were aligned to the *Z. tritici* IPO323 reference genome (58) using Bowtie2 v2.4.1 (74) with the option --very-sensitive-local. Duplicated sequences were tagged and removed using the Picard MarkDuplicates function v2.27.4 (http://broadinstitute.github.io/picard). ChIP-seq data quality was analyzed using the SSP (strand-shift profile) tool v1.1.0 (https://github.com/rnakato/SSP) with short background lenght option to quantify the signal-to-noise ratio (NSC), response signal correlation (RSC) and the mapped-read distribution throughout the genome (Bu). We used the Picard tool EstimateLibraryComplexity to calculate the library complexity (*i.e.* the non-redundant read fraction per 10 million mapped reads). The FRiP score (fraction of reads in peaks) was calculated using SAMtools v1.6 (75) to count total mapped reads and BEDTools v2.30.0 (76) to intersect and quantify reads overlapping peak regions. Peak calling was performed using the findPeaks function in the software package Homer (version 2.6.6) (http://homer.ucsd.edu/homer/ngs/peaks.html) with a peak calling size of 800 base pairs, which specifies the width of peaks that will form the basic building blocks of peaks in the regions (77). The minimum distance between adjacent peaks was set to 800 bp. ChIP-seq peaks were annotated for nearby gene features using BEDTools (version v2.30.0) (76), and gene models predicted for the IPO323 genome by Lapalu et al. (58) were used.

### Genomic window analyses

We analyzed genomic factors associated with previously generated H3K27me3, H3K4me2, H3K9me3 ChIP-seq datasets aligned to the IPO323 reference genome (54). We defined windows as non-overlapping 10 kb intervals using BEDTools v2.30.0 (76) and we calculated the percentage of base pairs covered by ChIP-seq peaks within each 10kb bin. For correlation analyses, the percentage of base pairs covering each bin was also calculated for other available parameters from the reference genome (IPO323): *i.e.* annotated genes (58); TEs detected using a curated *Z. tritici* TE consensus library (78); and recently inserted TEs (48);GC content per 10kb bin calculated using geecee v6.6.0.0 (https://www.bioinformatics.nl/cgi-bin/emboss/geecee). We also analyzed 3D genome organization features such as TADs (55) to assess H3K27m3 mark distribution. We considered H3K27m3 TAD coverage <20% in all isolates as no coverage and >20% as consistent coverage. Variable-switch was defined as isolates exhibiting TADs with extreme coverage variation (below 20% and above 80% coverage among isolates, respectively), whereas variable-nonswitch referred to isolates having TADs spanning the full H3K27m3 coverage spectrum, including at least one isolate with no coverage.

We further explored genomic features associated with H3K27m3 variation by ChIP-seq read coverage generated from the 45 field isolates. We used the same non-overlapping 10 kb intervals for summary statistics. To discard possible false negative ChIP-seq results caused by segmental deletions, we used previously generated copy number variation (CNV) data (79) obtained in intervals of 1kb using GATK CNV caller v4.1.9.0 (80). We considered genomic segment to be absent in a particular isolate if half or more of it was called as missing by the CNV calling step. For each isolate, we calculated the percentage of base pairs covered by H3K27m3 peaks per 10kb bin. We then computed the mean and coefficient of variation (CoV) across isolates to quantify average enrichment and variability. Then we used isolate-specific new TE insertion data and summarized evidence for non-reference insertions per 10kb bin (78). The number of non-reference TE insertions across all isolates per bin was summed and used as a parameter in correlation analyses.

### Population variation analyses

Individual gene loci were analyzed for evidence of gene deletions to remove erroneous calls of H3K27m3 peak variation caused by the lack of the underlying sequence in some isolates. A gene was considered deleted in a particular isolate if half or more contain null CNV calling, as described above. RNA sequencing data was accessed from Abraham et al. (42). The RNA-seq data was generated from isolates grown in modified Vogel’s Medium N (Minimal) replacing ammonium nitrate with potassium nitrate and ammonium phosphate. The media was devoid of any sucrose and agarose to induce hyphal growth. The NucleoSpin® RNA Plant and Fungi kit were used to extract total RNA from filtered mycelium after 10-15 days. An Illumina HiSeq 4000 was used to sequence TruSeq stranded mRNA libraries with 150 bp inserts in single-end mode. We checked RNA sequences for quality using FastQC v0.11.5 (72) and trimmed using Trimmomatic v0.36 (73) to remove adapter sequences and low-quality reads. We aligned trimmed sequences to the *Z. tritici* reference genome using HISAT2 v2.1.0 (81) with the parameter “--RNA-strandedness reverse”. The reads mapped to gene models were counted using the QTLtools v1.1 (82) in --Quan mode. Normalization of the counts was done using the –rpkm option implemented in QTLtools.

To address H3K27m3 gene coverage within the population, we considered H3K27m3 gene body coverage below 20% as no evidence for coverage, above 80% as full coverage, and intermediate percentages of overlap of H3K27m3 marks as partial coverage. Consistent full and lacking coverage was considered if at least 80% of the isolates exhibited the given pattern. The remaining were considered variable.

### Gene function enrichments

Effectors were predicted using EffectorP v3 (83) based on *Z. tritici* IPO323 reference genome (58). Secondary metabolites gene clusters were predicted using antiSMASH v.5.0 (84) also based on the reference genome.

## Declarations

### Author contributions

L.N.A. and D.C. conceived the study. L.N.A. and S.M.T performed experiments. L.N.A., A.M.S. and S.B. performed analyses. L.N.A., A.M.S. and D.C. wrote and revised the manuscript. A.M.S. and D.C. supervised the work.

### Data availability

ChIP-seq datasets are available from the NCBI Sequence Read Archive accession number PRJNA1321741.

## Acknowledgments

We are grateful for the helpful suggestions by Mareike Möller on protocols for chromatin immunoprecipitation.

