## Supplementary Figures for "Intra-population variability in genome-wide repressive histone marks underpins differential gene expression in a fungal wheat pathogen"

Abraham et al.

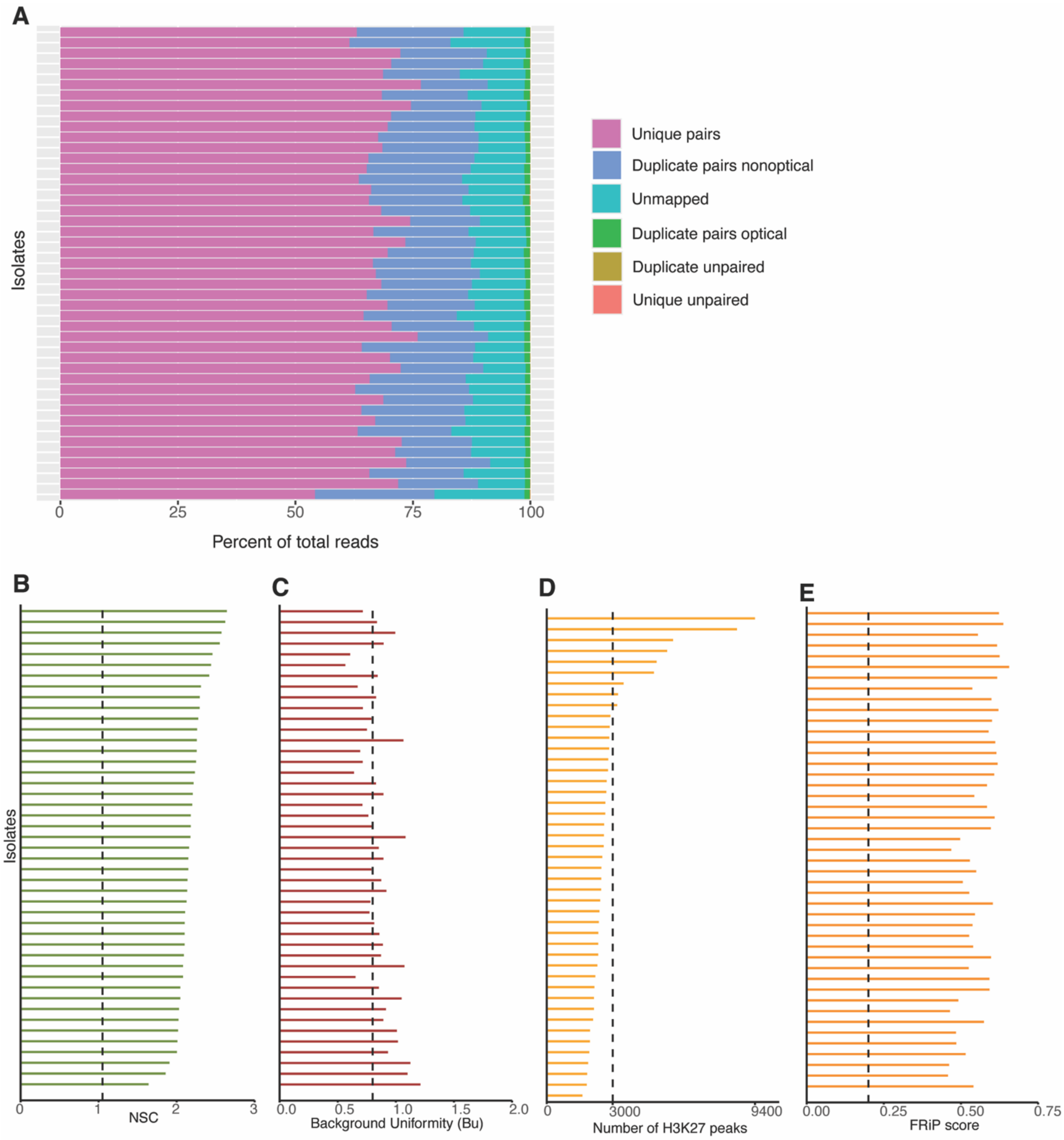


**Supplementary Figure S1:** A) Percentage of total reads mapped to the *Z. tritici* reference genome IPO323. B) Normalized strand cross-correlation coefficient (NSC) of aligned ChIP-seq reads. C) Background uniformity (Bu) of aligned ChIP-seq reads. D) Number of H3K27 peaks among isolates in the population. E) Fraction of reads in H3K27 peaks (FRiP score).

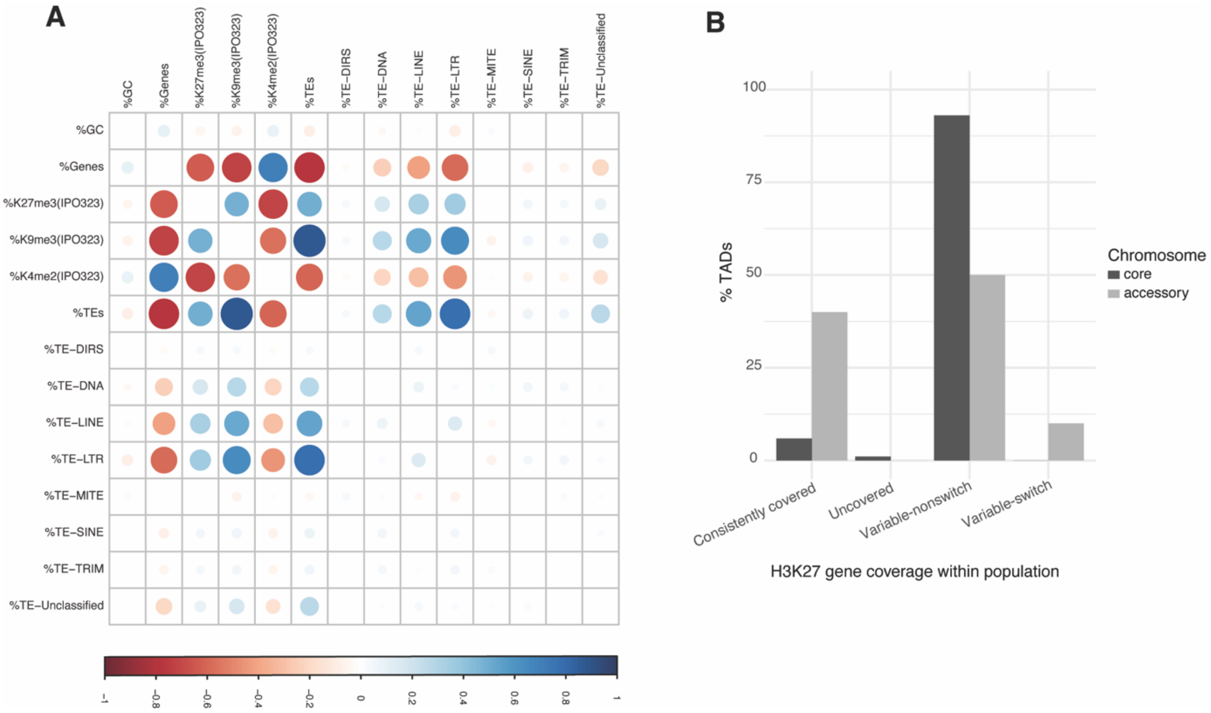


**Supplementary Figure S2:** A) Correlation plot between the percentage of 10-kb bins covered by GC, H3K27, H3K9, H3K4, genes, and TEs in IPO323 *Z. tritici* reference genome. B) Number of Topologically Associating Domains (TADs) in each H3K27 TAD coverage within population.

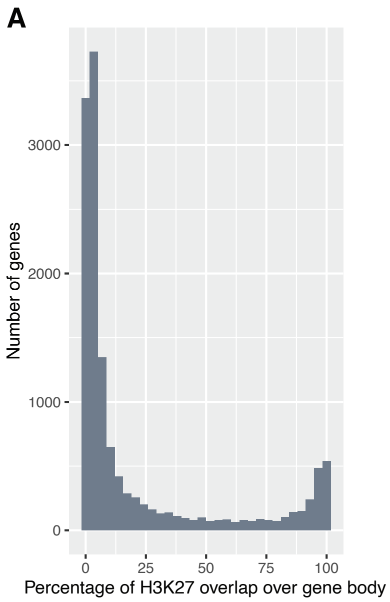


**Supplementary Figure S3:** A) Distribution of H3K27 mark coverage across genes.

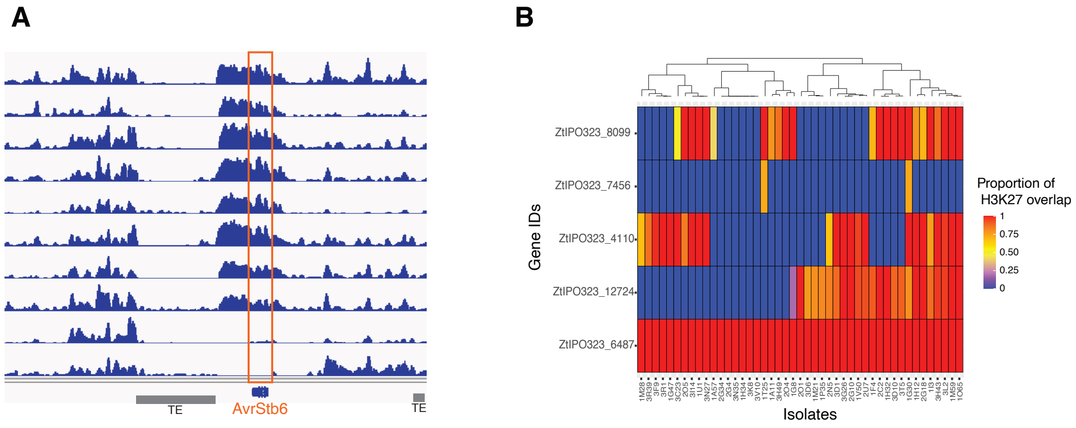


**Supplementary Figure S4:** A) H3K27m3 mark profile in *AvrStb6* gene among a subset of isolates in the population. B) H3K27 m3 coverage over candidate effector genes with variable expression over infection.
